# Dosimetric Characterization and Workflow Optimization of the FLASH-SARRP for Reliable Preclinical Radiobiological Studies

**DOI:** 10.64898/2026.07.06.736680

**Authors:** Michèle Knol, Patrik Gonçalves Jorge, Louis V. Kunz, Pierre Korysko, Benoît Petit, André Durham, Vozenin Marie-Catherine, Pelagia Tsoutsou, Nikolaos Koutsouvelis, Julie Lascaud

## Abstract

**Objective:** The reliable use of preclinical small-animal irradiators requires dosimetric characterization and quality assurance (QA) protocols. This study aimed to characterize the FLASH-SARRP, a dual-tube small-animal X-ray irradiator capable of ultra-high dose-rate (UHDR) delivery, as well as to establish and validate a practical QA workflow that improves dose reproducibility for preclinical FLASH experiments.

**Approach:** A combined dosimetric approach using Gafchromic films for spatial dose assessment and a plastic scintillating fiber for real-time temporal monitoring, was used to evaluate beam characteristics. Custom 3D-printed spacers were designed to ensure reproducible experimental configurations. The temporal pulse structure of the two X-ray tubes and their synchronization during simultaneous irradiation were investigated using scintillating fiber measurements. Based on the findings of the two dosimeters, a QA workflow was implemented and validated, which uses a cross-calibration between the Gafchromic film and the scintillating fiber in the experimental setup, followed by pre-irradiation verification with the fiber to compensate for day-to-day output variations.

**Main results:** Simultaneous irradiation of both X-ray tubes achieved dose-rates of approximately 80 Gy/s inside a homogeneous irradiation field suitable for small-animal studies. Temporal measurements revealed an average delay of 10 ms in between the two X-ray tubes, resulting in temporal dose-rate heterogeneity throughout the delivery of the dose. Additionally, a substantial inter-session variability of approximately 11% was found, whereas the intra-session variability was relatively low around ∼4%. Implementation of the proposed QA workflow reduced the inter-session variability to around 5%, approaching the intrinsic intra-session variability of the machine.

**Significance:** This work highlights the importance of temporal dosimetry for preclinical FLASH studies, as a measurable desynchronization between the two X-ray tubes was identified, generating temporal dose-rate heterogeneity that would not be detected by integrated dose measurements alone. The proposed QA framework improves reproducibility and provides a practical approach for reliable preclinical investigations.

## Introduction

FLASH radiotherapy is characterized by the delivery of radiation at ultra-high dose-rates (UHDR) and has been shown to reduce normal tissue toxicity while maintaining anti-tumor efficacy compared to radiotherapy at conventional dose-rates, a biological phenomenon called the FLASH effect (Favaudon *et al* 2014). The FLASH effect has now been demonstrated in various *in vivo* experimental models (Vozenin et al 2022, 2026) with several beam types, including electrons, protons, photons and carbon ions (Vozenin *et al* 2024). Today, the major challenges in FLASH research are related to accelerator technology, dosimetry and treatment planning system development, as well as to understanding the biological basis of the FLASH effect. Preclinical FLASH research combines both aspects, understanding the mechanisms of the differential effect in normal tissues versus tumors and defining the optimal irradiation parameters required to elicit the FLASH effect (Knol *et al* 2026).

Further research is needed to advance photon-FLASH beams, as photons are the most widely used type of radiation in the clinic, and therefore the most likely to be largely adopted by the radiation oncology community. Unfortunately, photon FLASH remains technologically the most challenging, as reaching ultra-high dose-rates with bremsstrahlung targets is a significant limitation due to target heating. Nevertheless, the technology is rapidly evolving, and photon-FLASH platforms are becoming available (Maxim *et al* 2019, Lin *et al* 2025). Experimental data are now also emerging from photon-FLASH systems. The first report of the FLASH sparing effect induced by photons in the brain was published already in 2018 using a synchrotron (Montay-Gruel et al 2018) and recently more biological results have been made available (Gao et al 2022, Miles et al 2023, Brown et al 2025, Lin et al 2025, Moreau et al 2025, Ghita-Pettigrew et al 2026, Zhang et al 2026, Hao et al 2026a) stimulated by the development of photon FLASH technological platforms at Hopkins by the group of Wang and Rezaee, now commercialized as the FLASH-SARRP (Xstrahl, Inc). This is a self-shielded cabinet equipped with two opposing kilovoltage X-ray tubes with rotating anodes, enabling both UHDR as well as conventional irradiations of small animals (Rezaee *et al* 2021, Miles *et al* 2023, 2024, Tajik Mansoury *et al* 2025). In parallel, the group of Zhang and Du (Gao *et al* 2022, Lin *et al* 2026, Hao *et al* 2026b) have also developed a 10 MV clinical photon-FLASH linac, called CHEX-FLASH, commercialized by Zhongjiu Flash Medical Technology Co.,Ltd, opening new opportunities for clinical translation.

In order to carry out meaningful preclinical and biological FLASH experiments, irradiation systems must not only be able to reach dose-rates sufficient to generate the FLASH effect, typically higher than at least 40 Gy/s but also deliver stable and reproducible irradiation conditions. To allow partial irradiation of small animals, a homogeneous irradiation field of at least 100 mm^2^ is needed. Moreover, the system should ensure reproducible dose delivery in subsequent irradiations, as well as from day-to-day, as even slight differences in the delivered dose, dose-rate, or pulse structure may influence biological outcomes and thereby compromise the interpretation and reproducibility of experimental results (Vozenin *et al* 2020, Böhlen *et al* 2024). Therefore, defining a robust quality assurance (QA) workflow is essential to ensure precise and reproducible biological experiments, which is the aim of this present work. Additionally, this QA workflow must account for the specific configuration of the FLASH-SARRP system, which includes two X-ray tubes. This introduces additional complexity in terms of temporal synchronization, geometric alignment between both tubes, and subsequently the homogeneity of the irradiation field.

The aim of this study was, therefore, to establish a QA workflow for two-tube X-ray systems applied to the FLASH-SARRP to support future preclinical and biological FLASH experiments under well-characterized and reproducible irradiation conditions. To this end, we implemented Gafchromic films (Ashland Inc.) and a plastic scintillating fiber (Luxium solutions) into custom-designed spacers to characterize the system under various experimental configurations and to account for the day-to-day variation of the system. While films are well-suited for dose measurements and offer a high spatial resolution (Jaccard *et al* 2018, Jorge *et al* 2019), plastic scintillating fibers have a very fast response time in the range of nanoseconds, a linear dose response, near water-equivalence, and can easily be integrated into an experimental setup thanks to their small size and flexibility (Hart *et al* 2022, Baikalov *et al* 2025). This workflow will support future preclinical and biological FLASH studies by providing a robust framework for determining acceptable variations in tube synchronization and system stability to sustain the FLASH effect.

## Methods

### Irradiation Setup

All irradiations were performed with the FLASH-SARRP system using either a single tube or both X-ray tubes simultaneously. In FLASH mode, the tube current has a range of 10-630 mA, and a peak voltage of 150 kVp, with selectable irradiation times between 1 and 250 ms in discrete increments. No external filtration was applied; however, an internal filtration of 0.7 mm aluminium was present within the tubes. The nominal half-value layer (HVL) of the beam is 3.7 mm Al. External fans were used between irradiations to enhance cooling of the machine and prevent overheating due to the fast rise in temperature of the tubes. Depending on the experimental setup, such cooling measures may be required to maintain the tube temperatures within the operational range and should be considered when designing and operating an X-ray UHDR irradiation system. The distance between the tubes was minimized (i.e., 7 cm distance between irradiation center and the tube sources) to reach the highest dose-rates achievable with this system.

A set of dedicated 3D-printed spacers was designed to ensure reproducible positioning of the tubes, samples and dosimetry tools across experiments. The base structure of the 3D-printed platform used for the dosimetry experiments is depicted in Fig. 1a-c. The two open extremities are shaped to fit to the tube surface, while an intermediate layer is printed at mid-height with a 6 x 6 mm^2^ central aperture to insert the Gafchromic film and a transverse through-hole to guide the scintillating fiber. The design of the 3D spacer was optimized to minimize interactions with the primary beam. This baseline platform was adapted to fit the needs of future biological experiments, including irradiations of biological samples (e.g., zebrafish embryos) in a Petri dish with both tubes (Fig. 1e), or irradiations of the legs of small animals using only the bottom tube (Fig. 1d). Extra height adjusters were designed to be placed between the printed platform and the tube to enable precise modification of the source-to-surface distance (SSD). All elements were modelled in the open-source Tinkercad software and printed by a BambuLab A1 mini, using either PLA or PETG filaments. A 0.4 mm nozzle, 0.2 mm layer thickness and a standard infill of 15% were used.

**Figure 1.**
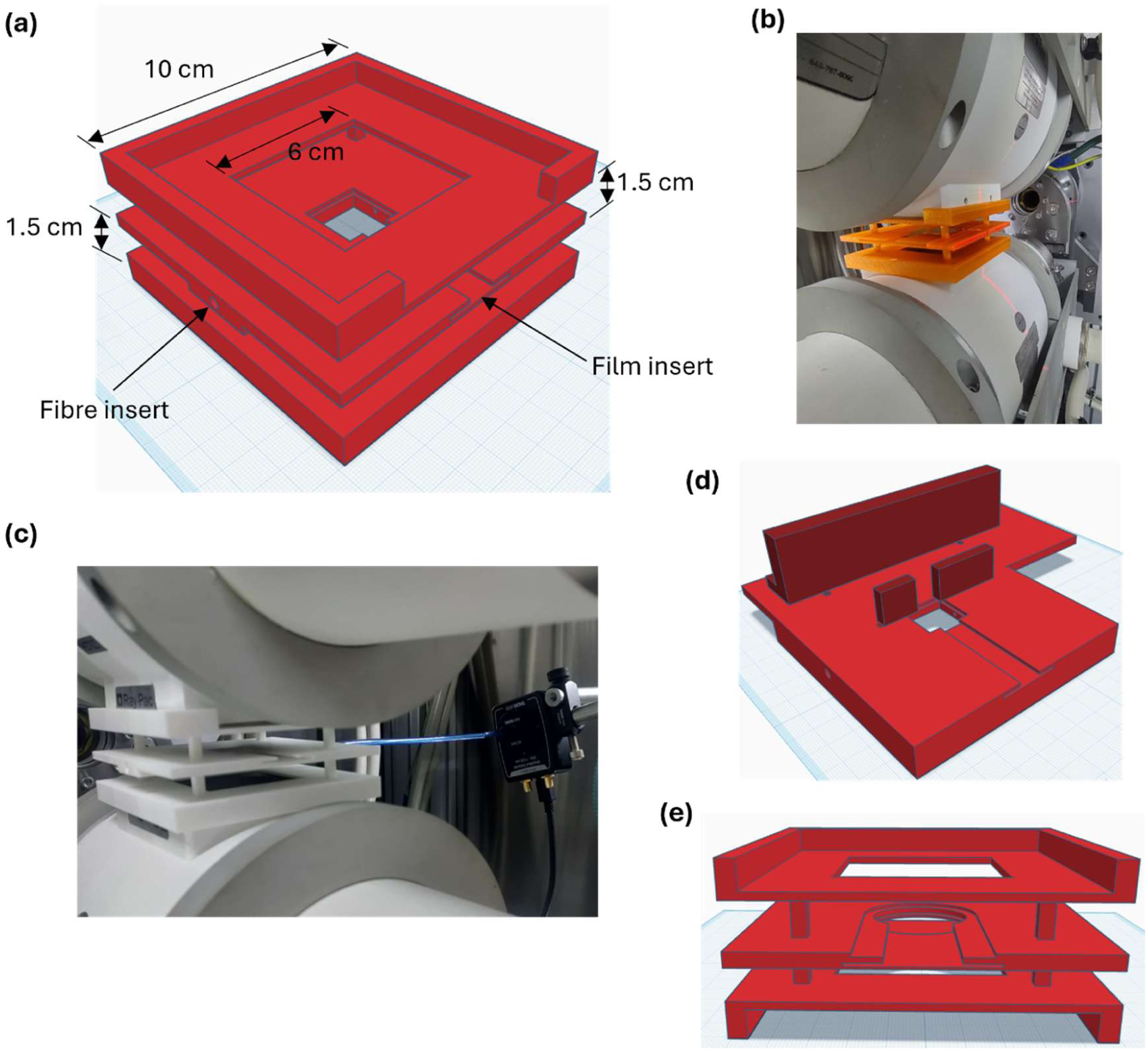
The setup of the film and fibre measurements in the SARRP FLASH. **(a)** 3D model of the spacer including a slit for Gafchromic positioned at a distance of 7 cm from the source of both tubes and a hole for the plastic scintillating fibre directly below the film. Model created in Tinkercad. **(b)** A photo of the spacer setup between the two tubes used for film measurements. **(c)** A photo of the spacer setup with the plastic scintillating fibre and connected to the photodetector. **(d)** 3D model of the spacer used for knee irradiations using only the bottom tube. A slit for Gafchromic film and a hole for the plastic scintillator is implemented in a similar way to the original spacer in A. **(e)** 3D model of the spacer used for Zebrafish embryo irradiations with both tubes.

### Film Dosimetry

To assess the shape and uniformity of the beam, as well as to obtain precise dose measurements, EBT-XD Gafchromic film was used. The films were calibrated against the SRT-100 (Sensus Healthcare, United States) at the Geneva University Hospitals (HUG), a 100 kV clinical photon irradiator calibrated and traceable to primary standards at the Swiss institute of metrology (METAS). To assess the validity of the calibration curve over time, a verification was performed every two months by remeasuring selected dose points and comparing them to the original calibration. After 24 hours of development in a dark environment, films were scanned with an Epson Expression 10000XL flatbed scanner, using the positive film mode at 300 dpi without further corrections and saved as a tiff file. Files were analyzed with a custom Python script, using the red channel. An unirradiated film that underwent the same process as the irradiated films was used to determine the reference signal *I*_0_. The net optical density (netOD) was calculated as 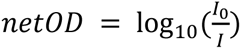, where *I* is the signal measured in the irradiated film. netOD values were subsequently converted to dose using the calibration curve. The overall uncertainty of the film dosimetry was determined to be 4%, including contributions from the calibration fit, film reproducibility, scanner reproducibility, and image analysis.

To independently validate the film dosimetry procedure, dose measurements obtained with the EBT-XD films were compared with measurements from a soft X-ray ionization chamber (TN23342, PTW Freiburg, Germany). Single-tube irradiations were performed at exposure times ranging from 32 and 250 ms to cover the dose range of the FLASH-SARRP. Ionization chamber and film measurements were acquired sequentially using the same irradiation setup during the same experimental session. Both film and ionization chamber were calibrated using the SRT100. A linear regression to assess the agreement between both methods demonstrated excellent agreement between both methods with a slope of 1.0058 and R^2^ = 0.9996 (Supp. Fig. 2a,b).

When both tubes were used, the reported dose was defined as the average value in the homogeneous central part of the field within a circular region of interest (ROI) of 10 mm diameter. When only the bottom tube was used, the homogeneous part of the field was located towards the top of the field, where a circular ROI of 10 mm was used to determine the average dose.

### Depth dose measurement, linearity and field homogeneity assessment

Percentage Depth Dose (PDD) curves of the beam central axis were obtained using 1 cm-thick RW3 solid water slabs (PTW-Freiburg GmbH, Freiburg, Germany) (50×50 mm) stacked between both tubes (Supp. Fig. 4a). Gafchromic films were placed between the solid water slabs and at the surface of both tubes. Irradiations were performed using identical parameters (150 kV, 630 mA, 100 ms) with either one or both tubes.

Dose linearity with exposure time and tube current were evaluated using the spacer from Fig. 1a and Gafchromic film. Films were irradiated subsequently by varying either time or current, while keeping the other parameters constant (150 kV, 630 mA, 100 ms).

The homogeneity of the irradiation field at the foreseen biological sample position (i.e., at intermediate distance between the tubes) was evaluated by irradiating Gafchromic films with the dosimetry spacer shown in Fig. 1a. The dose distribution within the 10 mm circular ROI in the middle of the field was verified on six films that were irradiated during different sessions. Homogeneity was quantified as the uniformity inside the ROI, defined as the standard deviation of the dose normalized to the mean dose: 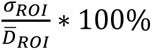.

Flatness was calculated within the central 80% of the field, defined by the 50% isodose lines. It was defined as the percentage variation between the minimum and maximum dose within this region: (*D*_max_ − *D*_min_)⁄(*D*_max_ + *D*_min_) ∗ 100%. Symmetry was evaluated over the same region and was defined as the maximum percentage difference between doses measured at points equidistant from the central axis: *max*(V *D*(*x*) − *D*(−*x*) V)⁄*D*_centre_ ∗ 100%.

Reproducibility was assessed across eight independent sessions spread over several weeks. Both tubes were used at 150 kV, 630 mA, and for a total irradiation time of 125 ms to target a dose of about 10 Gy. To evaluate the reproducibility, film dose distributions were compared directly using film-to-film gamma analysis. Gamma index analysis was performed using criteria of 2 %/2 mm, adapted to the small scale of preclinical irradiations. Comparisons were performed both between repeated irradiations during the same session and between irradiations performed during different sessions. For each comparison, one film was used as the reference distribution and compared with a second film. Three independent film pairs were analyzed for each category.

### Fiber measurements

To obtain information about the dose variation over time at the pulse scale, a plastic scintillating fiber (BCF-60XL, 1 mm diameter, Luxium solutions) was used. The fiber was mounted to a silicon amplified photodetector with an integrated acquisition system (PDA36AU, Thorlabs). The fiber output signal was amplified by 70 dB and acquired using the provided PDA_USB software at a sampling rate of 1 kHz.

Fiber measurements were analyzed using a custom Python script. The signal was filtered in post-processing by using a moving average and subsequently a zero-phase Butterworth low-pass filter of the fourth order. The start and the end of the pulse were detected by their rising and falling edges using an adaptive threshold set to 20% above the baseline. From this, the total irradiation time was determined, as well as the delay between the tubes, the amplitude, background noise, and the area under the curve (AUC).

Reproducibility measurements were taken during various sessions with only the fiber integrated in the two-tube dosimetry spacer (Fig. 1a), always using the same irradiation parameters (150 kV, 630 mA, 100 ms). Only sessions acquired without modification of the fiber-photodetector configuration were included in the analysis.

### QA workflow

The proposed QA workflow is divided into two steps: (1) cross-calibration of the scintillating fiber against film dosimetry and (2) pre-experiment verification of the delivered dose-rate. As the fiber provides a signal that is proportional to the dose output, but does not directly measure the dose, a cross-calibration against film dosimetry was performed. Calibration measurements were carried out on the days preceding the experiments of interest (i.e., biological experiments), using the exact irradiation setup; the spacer was not moved between the calibration and the experiment. During the calibration, the fiber and films were irradiated simultaneously using the same voltage and current as for the experiment. Typically, five irradiations were performed while varying the irradiation times between 32 and 250 ms. Afterwards, the relationship between the AUC from the fiber signal and the measured dose from the films was determined, resulting in a cross-calibration factor between the AUC and absolute dose.

Next, on the day of the validation experiments, several irradiations were performed after the system warm-up. For each irradiation, the fiber signal was used to determine an instantaneous dose-rate surrogates (AUC/ms) from the average peak power and the pulse duration. Using the previously established calibration factor, these values were converted to instantaneous dose-rates (Gy/s). In case of large deviations, the source-to-surface distance (SSD) and/or the pulse duration was adapted to match the requested dose-rate and dose, assuming an inverse square relationship between the deposited dose and its distance from the source. To verify the validity of the inverse square approximation, additional film measurements were performed at varying SSDs using 3D-printed adjusters that were put in between the spacer and the tube to increase the distance of the sample to the source in a reproducible way. The measured dose-rate dependence was found to be consistent with inverse square scaling in the investigated range, considering the uncertainty attributed to set-up and positioning errors when changing the SSD (Supp. Fig. 3).

The robustness of the QA workflow was assessed for single-tube irradiation using the bottom tube and dedicated platform (Fig 1d). Several SSDs were tested in a single session using the height adjusters and multiple irradiation times between 50 and 250 ms were used.

### Statistical Analysis

The intra-session variability was calculated from the reproducibility film measurements. The coefficient of variation (CV) was calculated for each session as the ratio of the standard deviation to the mean of the measured doses. A pooled intra-session CV was then derived from a one-way ANOVA. Inter-session variability was calculated as the standard deviation of the session means divided by the overall average dose. For the validation of the QA workflow, doses were first normalized to the used irradiation time before determining the variation.

## Results

### Spatial dose distribution

The two-dimensional dose depositions obtained when using either the top tube only, the bottom tube only, or both tubes together, are depicted in Fig. 2a, with corresponding dose profiles extracted from the same Gafchromic film measurements in Fig. 2b. For single tube irradiation, profiles were heterogeneous, resulting from the heel effect, going in opposite directions for the two opposing tubes. The most homogeneous region of the field of the top tube is located towards the bottom of the field, while the bottom tube produced a homogeneous dose in the upper portion. When both tubes were operated simultaneously, their individual contributions complement one another, resulting in a centrally homogeneous dose distribution (Fig. 2a, b). Analysis of this central 10 mm diameter ROI used for dose reporting showed an average uniformity of 1.33 ± 0.10%, supporting the use of this ROI as a homogeneous dose distribution. Analysis of these dose maps showed an average field size of 4.2 x 4.0 cm, defined by the 50% isodose lines (Table 1). Furthermore, flatness was 9 % horizontally and 14 % vertically, consistent with the vertical asymmetry introduced by the heel effect, while symmetry was comparable across both directions (7%).

**Figure 2.**
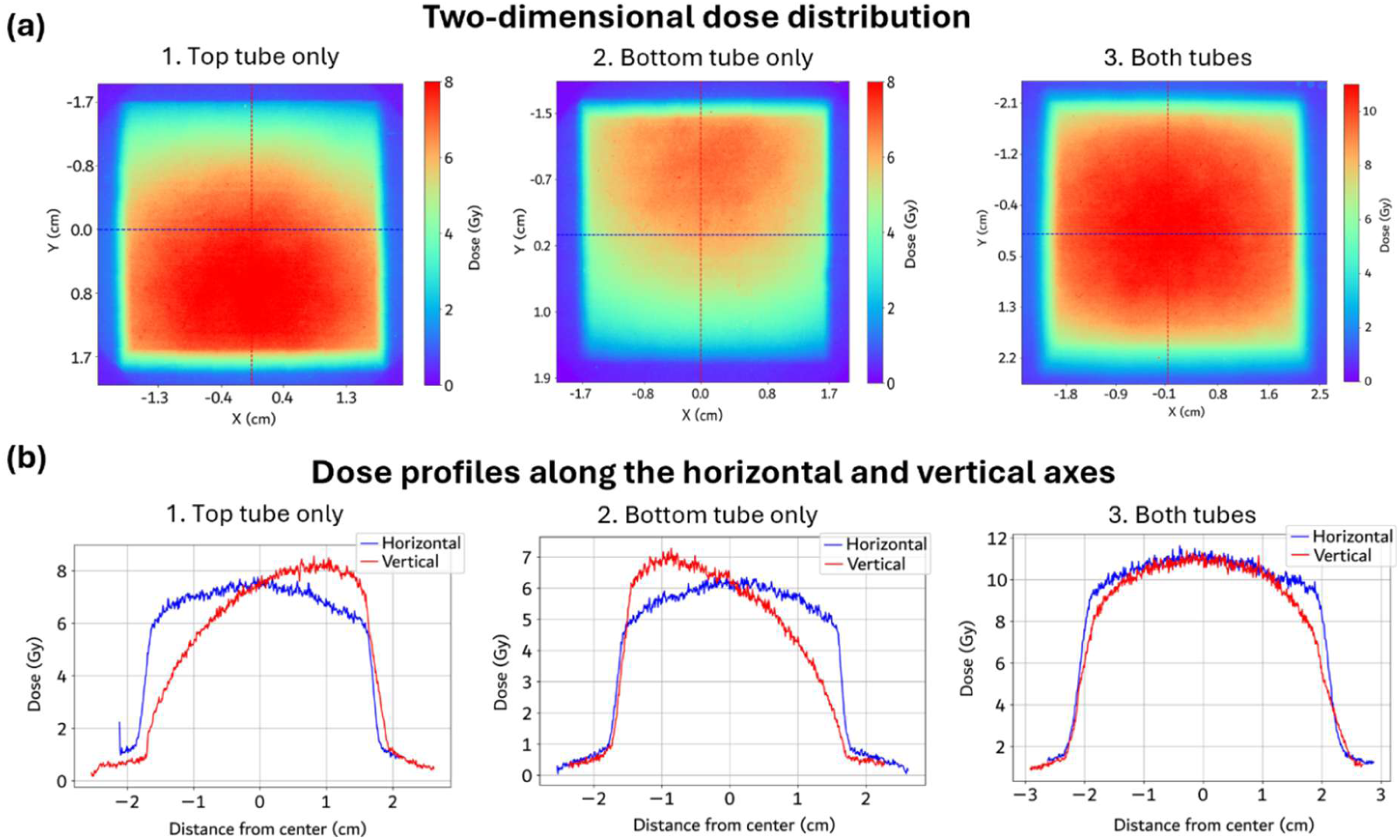
Beam characteristics. **(a)** Dose maps obtained from irradiated Gafchromic film with the top tube only (1), bottom tube only (2), or both tubes (3). The crosshairs through the middle of the field indicate the line used for the dose profiles. **(b)** Dose profiles through the middle of the field from the film irradiated with the top tube only (left), bottom tube only (middle), and both tubes simultaneously (right). The blue line indicates the horizontal line profile, the red line the vertical profile.

**Table 1.** Beam characteristics. Flatness was calculated as (*D*_max_ − *D*_min_)⁄(*D*_max_ + *D*_min_) ∗ 100% with the central 80% of the field (defined by the 50% isodose lines). Similarly, symmetry was calculated as *max*(V *D*(*x*) − *D*(−*x*) V)⁄*D*_centre_ ∗ 100%. The mean of 6 different measurements was taken in both the horizontal and the vertical direction. The field width was determined as the size between the 50% isodose lines.

| *μ* ± *σ* | **Flatness (%)** | **Symmetry (%)** | **Field width (cm)** |
|---|---|---|---|
| **Horizontal** | 10.6 ± 1.3 | 7.5 ± 0.7 | 4.19 ± 0.02 |
| **Vertical** | 16.3 ± 3.0 | 12.5 ± 4.9 | 3.99 ± 0.04 |

Depth dose measurement in FLASH-mode revealed a half-value layer (HVL) of around 14 mm water equivalent thickness. When both tubes were operated simultaneously, the dose distribution is homogeneous along the beam direction, with a slightly larger contribution of the top tube.

### Time profiles and tube synchronicity

The time profile of the X-ray pulse for a single-tube irradiation is presented in Fig. 3a for five deliveries using the bottom tube within the same session. As seen, the fiber output resembles a step-like function with a ramp-up at the start. The pulse structure within the session was highly reproducible in terms of both the duration and amplitude, and therefore, the AUC.

**Figure 3.**
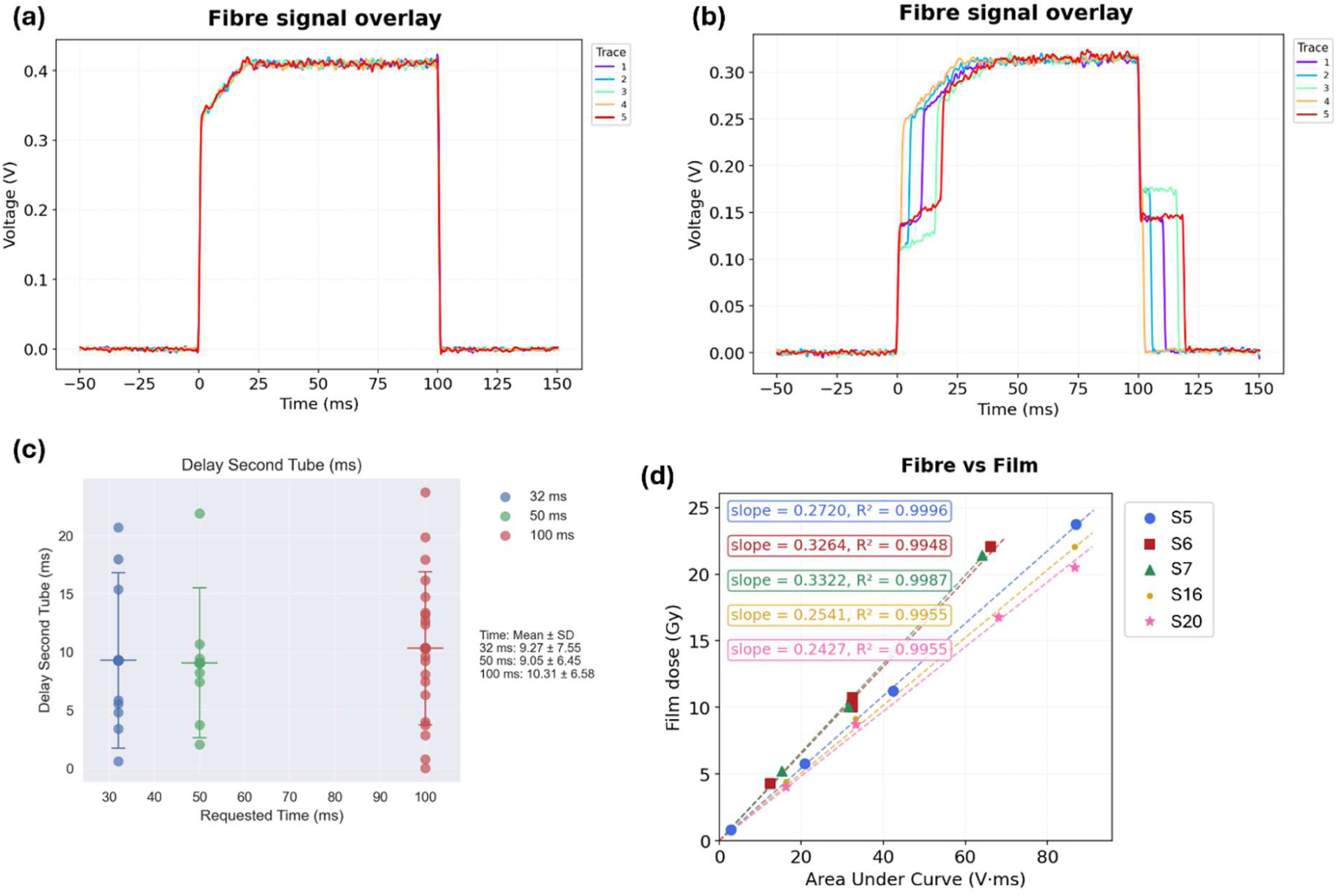
Fibre measurements of the delivery time profile. **(a)** Overlay of 8 irradiations measured during the same session with the same parameters. The smoothed signal is plotted. **(b)** Overlay of 5 irradiations (bottom tube only) measured during the same session with the same parameters. The smoothed signal is plotted. **(c)** The delay of the second tube, measured by the fibre measurements, defined as the time between the start of the first tube and the start of the second tube. Irradiations were performed during various sessions using either 32, 50, or 100 ms. **(d)** The area under the curve (AUC) of the fibre measurements and corresponding dose determined from Gafchromic films, together with a linear fit. The sessions during which the measurements were taken are listed in Supp. Table 1. The individual points represent single film measurements with an overall uncertainty of the film analysis of 4%.

When both tubes were operated simultaneously, fiber measurements revealed a systematic offset between the start of the irradiation of the two tubes (Fig. 3b). Although the overall shape of the two individual X-ray pulses and their requested irradiation time were preserved, they did not fire synchronously. The delay between the onset of the first and the second tube randomly varied between 0 and 24 ms, with a mean delay of 10 ms (Fig. 3c). The delay proved to be independent of the set irradiation time and no consistent pattern was observed as to which tube fired first. Consequently, the total irradiation time was increased, and the instantaneous dose-rate was reduced by a factor of two at the beginning and end of the pulse, introducing temporal dose-rate heterogeneity within a single irradiation delivery. This means that for a mean delay of 10 ms on a 100 ms irradiation this corresponds to a 10% increase in total irradiation time. This affects the overall dose-rate profile and, consequently, the instantaneous dose-rate, as the first and last 9% of the irradiation will be carried out with half of the dose-rate. The desynchronization is an even bigger concern for lower doses, since the delay proved to be independent of the total irradiation time. An average delay of 10 ms on a 32 ms irradiation (corresponding to approximately 3 Gy) will cause a 30% increase of the total irradiation time, as well as the first and last 25% of the irradiation with only half of the dose-rate.

### Intra- and inter-session variability

Intra- and inter-session variability were assessed by repeated measurements across multiple sessions. The results of the variability assessment are presented in Fig 4. The dose in the homogeneous part of the irradiation field varied between 8.3 and 12.4 Gy during the sessions, with a mean and standard deviation of 10.4±1.2 Gy (Fig. 4a), corresponding to an inter-session variability of 11%. The intra-session variability (4%) was slightly lower for the investigated parameters. This gave rise to dose-rates ranging across sessions from 66 to 99 Gy/s, with an overall average value of 82.8 Gy/s, whereas dose-rate variation within a single session was substantially lower (Fig. 4b).

**Figure 4.**
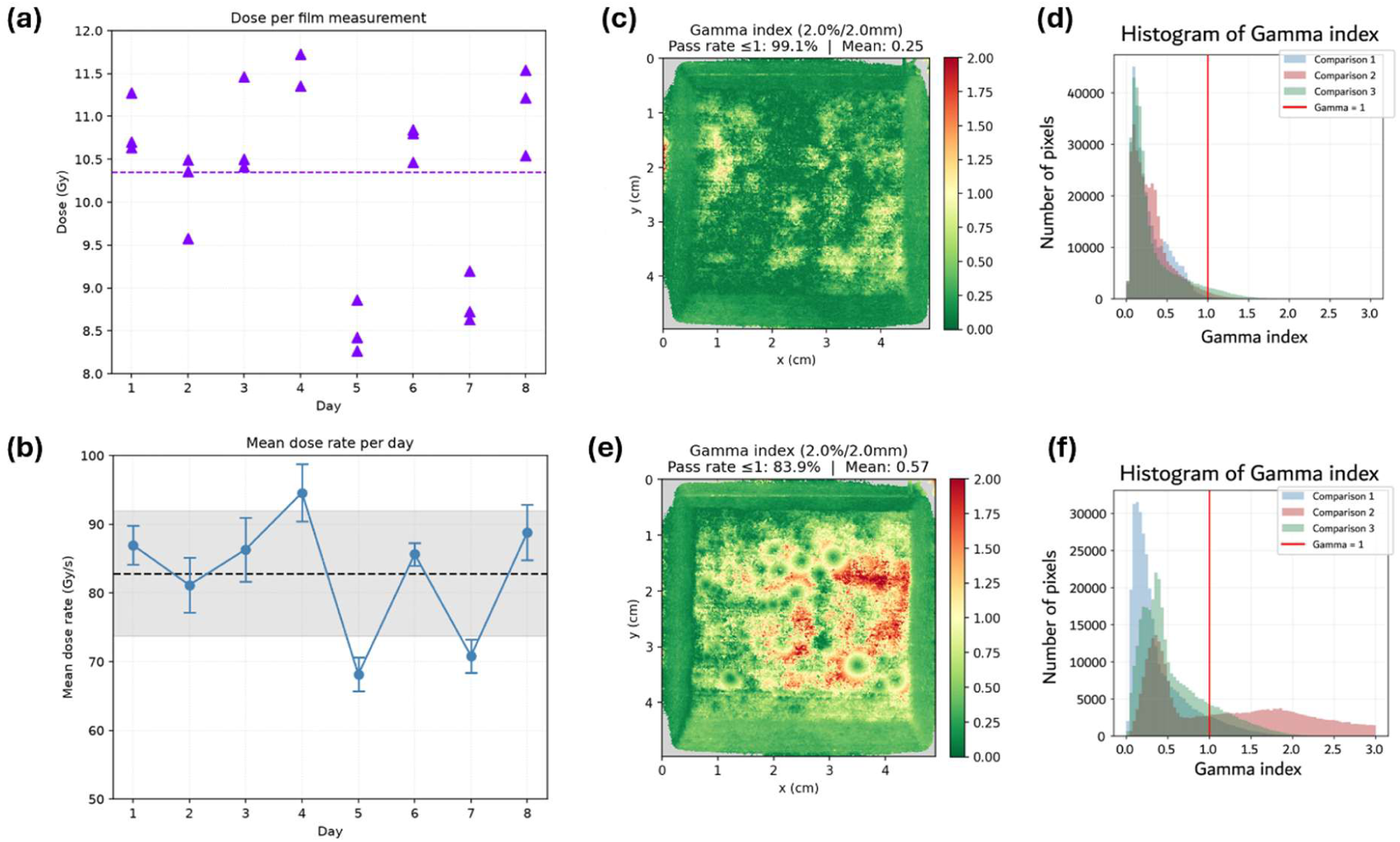
Intra- and inter-session variation of the irradiation field. **(a)** Measurements over several irradiation sessions with both tubes (Supp. Table 1). Dose was determined with Gafchromic film. Irradiations were all carried out with the same parameters (150 kV, 630 mA, 125 ms). The mean of all these measurements is shown with the dashed line (125 ms: 10.4 Gy). The individual points represent single film measurements with an overall uncertainty of the film analysis of 4%. **(b)** The dose rate over time determined with the dose from the Gafchromic film and the irradiation time from the measurements from (a). The mean of all measurements is shown with the dashed line (83 Gy/s), the grey area depicts one standard deviation. n = 3 per session. **(c)** A representative gamma index map (2%/2mm) between dose maps of irradiations performed during the same session (Supp. Table 1). **(d)** An overlay of histograms of the gamma indices of three comparisons of irradiations performed during the same session. **(E)** A representative gamma index map (2%/2mm) between dose maps of irradiations performed during two different sessions (Supp. Table 1) with the same parameters. **(f)** An overlay of histograms of the gamma indices of three comparisons of irradiations performed during three different sessions (Supp. Table 1).

The gamma index, which simultaneously accounts for spatial and dosimetric differences between irradiations, was used to evaluate the reproducibility of the dose distribution and field geometry. A gamma index below 1 is considered to be acceptable. For intra-session comparisons, a representative gamma index map demonstrated high agreement between irradiations, with a mean gamma index over the whole field of 0.25 and 99.1% of pixels passing the gamma criterion (Fig. 4c). The overlay of multiple intra-session comparisons confirmed this low variability (Fig. 4d). However, for inter-session comparisons a higher mean gamma index over the field of 0.57 was found, with only 83.9% of the pixels passing the threshold (Fig. 4e). Additionally, the overlay of several inter-session comparisons confirmed the substantially larger spread in both the dose magnitude and the field geometry compared to the intra-session results (Fig. 4f). Together, these results indicate significant day-to-day variations of the maximum dose and uniformity of the field that should be accounted for prior to biological experiments to ensure precise and reproducible irradiations.

The reproducibility was also assessed through fiber measurements. Irradiations were carried out over several sessions, using the same irradiation parameters. Overlaying the fiber signals from different sessions revealed a variability in the amplitude of the signal that was not observed during the intra-session comparisons of the fiber measurements, pointing again to the session-dependent output variation and showing that this is reflected in both film and fiber measurements (Fig. 5a). Quantification of the signal by determining the AUC across all fiber measurements confirmed a low intra-session variability of 0.25% and a substantially higher inter-session variability of 1.77% (Fig. 5b).

**Figure 5.**
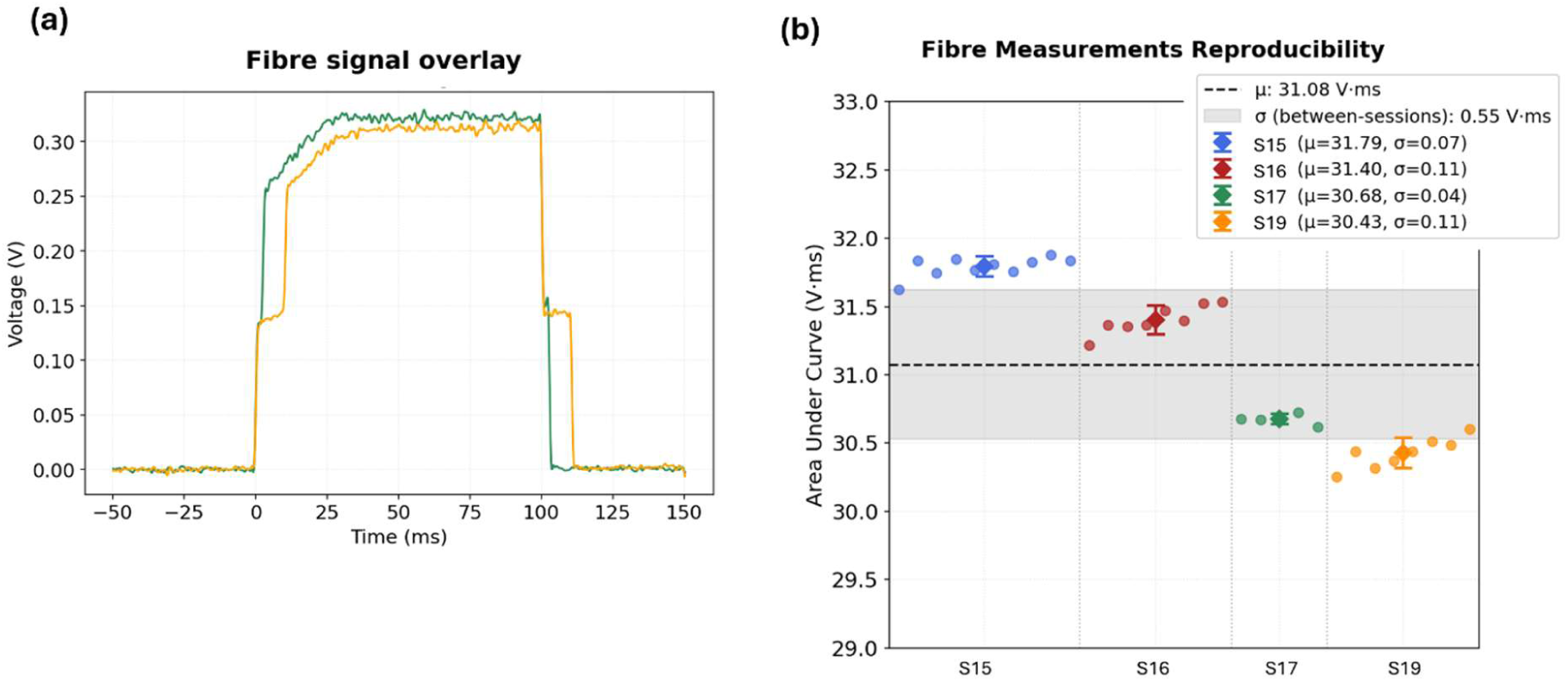
Evaluation of the fibre measurement reproducibility. **(a)** The smoothed signal of 2 measurements (100 ms) taken during different irradiation sessions (Supp. Table 1). **(b)** Integral of the fibre signal (area under the curve, AUC) taken during different irradiation sessions (n = 5-10). All measurements were taken during 100 ms irradiations using both tubes. The mean AUC of all measurements is depicted with the dashed line; the grey area shows 1 standard deviation. The mean and standard deviation are also shown per session (Supp. Table 1).

### Cross-calibration of the scintillating fiber and Gafchromic film

To establish a quantitative relationship between the scintillating fiber signal and the dose measured with Gafchromic films, simultaneous irradiations of film and the fiber were performed across multiple sessions using both tubes, while varying the exposure time. For each irradiation, the AUC of the scintillator signal was compared with the dose measured from the corresponding film, generating a calibration curve. A linear relationship was observed between the fiber AUC and the film-measured dose within each session (Fig. 6c), demonstrating that the response of the fiber was proportional to the delivered dose. However, it should be noted that the slope of the obtained calibration curves varied slightly between sessions, indicating that a session-specific calibration factor was required to convert the fiber signal into dose.

**Figure 6.**
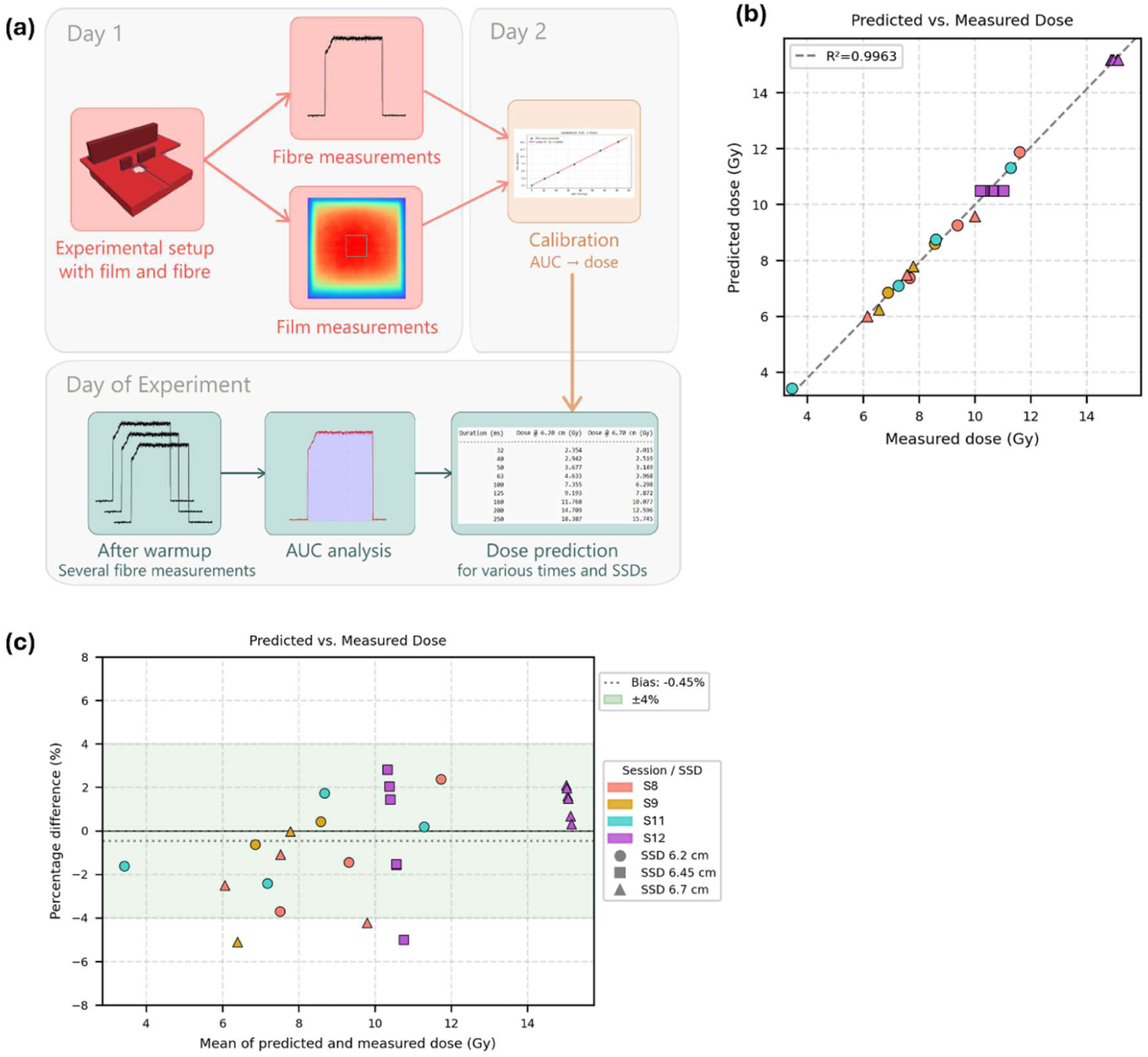
QA framework biological experiments. **(a)** Schematic of the workflow used for biological experiments. **(b)** Predicted versus the measured dose with a linear fit. Doses were measured with Gafchromic films over four different sessions. Doses were predicted according to the workflow presented in a. Measurements were performed using the spacer for knee irradiations as shown in fig. 1d. Colours represent the different sessions; shapes correspond to different SSDs (legend in c). The individual points represent single film measurements with an overall uncertainty of the film analysis of 4%. **(c)** The mean of the predicted and measured dose versus the percentage difference. Measurements are the same as in b. The inter-session variability is depicted as the green area. The individual points represent single film measurements with an overall uncertainty of the film analysis of 4%.

### Assessment of the QA workflow robustness

As the observed day-to-day variation of the FLASH-SARPP reached 11%, accurate dose delivery for preclinical and biological FLASH experiments cannot rely only on fixed irradiation parameters for a given targeted dose. Therefore, a QA workflow was developed to account for this variation by establishing a calibration between scintillating fiber measurements and film dosimetry, followed by a verification of the output of the machine just before sample irradiation and subsequent adjustment of the irradiation time and/or the SSD when necessary (Fig. 6a).

The performance of this workflow was validated using the bottom-tube irradiation setup and the 3D-printed spacer from Fig. 1d. Predicted doses obtained from the QA workflow were in good accordance with the measured doses from the films, showing a linear relationship with an R^2^ of 0.9963 (Fig. 6b). These irradiations were performed using various irradiation times and SSD, thus validating a large range of the predicted doses. The percentage difference between the predicted and the measured dose showed a small average bias of −0.45% (Fig. 6c). The resulting variation between days when the workflow was used was 4.7%, substantially lower than the intrinsic day-to-day variation observed in the system output. These results demonstrate that the QA workflow can compensate for a large fraction of the output variability and improve dosimetric reproducibility for preclinical and biological experiments.

## Discussion

In this work, we characterized the temporal and spatial dose delivery of a photon-FLASH platform by combining the use of a plastic scintillating fiber as a real-time monitoring tool with Gafchromic films for absolute dosimetry. Gafchromic films revealed a dose distribution, field size, and depth dose curve, consistent with previously reported results from similar FLASH-SARRP systems (Tajik Mansoury *et al* 2025, Miles *et al* 2024). Meanwhile, the flatness and symmetry values exceeded those typically accepted for clinical irradiators, and they are also slightly higher (7%) than comparable preclinical irradiation systems (∼ 3%) (Wang et al 2017, Feddersen et al 2019). However, a homogeneous circle of 1-cm diameter in the center of the field is sufficient for the irradiation of small animals. Together, these results provide a comprehensive spatial characterization of the beam, which is crucial for establishing accurate dosimetry for future preclinical studies.

The implementation of a scintillating fiber in the FLASH-SARRP setup revealed a substantial desynchronization between the two tubes, with an average delay of 10 ms. Such a delay extends the effective total irradiation time beyond the requested total irradiation time. Critically, this heterogeneity of the instantaneous dose-rates within one irradiation can influence whether the FLASH effect is reliably achieved (Liu *et al* 2025, Zhu *et al* 2025). With an average dose-rate of around 80 Gy/s, the time during which only one tube is irradiating corresponds to dose-rates of around 40 Gy/s, a value that might not be sufficient to trigger the FLASH effect. Additionally, since the delay varies substantially, the extent of the change in instantaneous dose-rate will vary from irradiation to irradiation and will thus not be reproducible. This poses an extra problem when fractionated regimens are used, typically with lower doses per irradiation, since not all fractions will be delivered with the same dose-rate, possibly leading to higher chances of losing the sparing effect, and without regulation of the FLASH delivery. At present, the FLASH sparing effect with the FLASH SARRP has only been reported *in vivo* using one tube (Brown *et al* 2025, Ghita-Pettigrew *et al* 2026). *In vivo* studies remain to be done using both tubes to see if a sparing effect can reliably be found. In the meantime, a solution needs to be found to significantly decrease the offset.

Beyond the technical implications, the finding of the desynchronization of the two X-ray tubes also highlights the importance of independent characterization of preclinical irradiation platforms. Unlike clinical machines, preclinical irradiators are often operated mainly by biologists and may not always undergo extensive commissioning or routine QA protocols by medical physicists. Under these circumstances, users may reasonably assume that the programmed irradiation time accurately reflects the delivered irradiation time. Our findings demonstrate that such assumptions might not always hold and more importantly, may affect both the reproducibility of the experiment and the interpretation of the biological outcomes. This observation emphasizes the need for standardized dosimetric procedures and the use of QA protocols in preclinical radiobiology research, as discussed in several other publications (Desrosiers *et al* 2013, Kirsch *et al* 2018, Hill *et al* 2025).

Comparing the output of the film to that of the fiber confirmed that both detectors capture the same underlying dosimetric variation and support the use of the fiber as a reliable real-time dosimetric monitor, complementary to film-based reference measurements. However, a variation was found in the calibration factor between the two dosimeters. This is attributed to variations in the repositioning of the fiber when remounting the irradiation holder between sessions (e.g., between sessions with one- or two-tube spacer) and is mitigated by systematically performing a new calibration after changing the irradiation platform. This weekly calibration was therefore implemented in the proposed QA workflow.

Both film and fiber measurements demonstrated an acceptable intra-session variability but consistently identified a higher inter-session variability. This increased inter-session variability complicates both the reproducibility and the standardization of FLASH delivery, a prerequisite when assessing subtle biological differences. Importantly, however, the real-time output of the plastic scintillating fiber enables adaptive delivery strategies, whereby the SSD or irradiation time can be adjusted to compensate for the day-to-day output variation, a correction that film-based dosimetry alone cannot support.

This work shows that the integration of a scintillating fiber into the FLASH-SARRP experimental setup can be used as both a characterization and an active QA tool for preclinical FLASH experiments. The characterization indicated significant day-to-day variations of the maximum dose and uniformity of the field that should be accounted for prior to biological experiments, to ensure precise and reproducible irradiations. To this manner, using the fiber as a QA tool can reduce the day-to-day variation to a similar level as the intra-session variation, while providing additional information on the time structure of the beam.

There are, however, several limitations to the proposed film- and scintillator-based dosimetry tool. First, the film calibration was established using the SRT-100 reference irradiator, which differs from the FLASH-SARRP in beam quality and HVL. Consequently, a systematic uncertainty associated with this indirect calibration cannot be completely excluded. Nevertheless, good agreement was observed between film-derived doses and ionization chamber measurements, suggesting that any resulting bias is small relative to the overall dosimetric uncertainty of the system. Furthermore, the scintillation yield has been shown to be temperature-sensitive, with changes between 0.5 and 3% per 5 degrees Celsius of heating (Buranurak *et al* 2013). Day-to-day variation of the room temperature or heating of the X-ray tubes during long series of irradiation could then directly impact the dosimetry outcomes if not taken into account. The proposed setup would therefore benefit from the integration of a temperature monitoring system for real-time temperature compensation.

A further limitation of the current setup is the sensitivity of the detected scintillation light to fiber positioning. To ease the mounting of the scintillator and irradiation platform within the FLASH-SARRP cabinet, the fiber must be unmounted every time, and no solid coupling was used between the fiber and the photodetector. Hence, small changes in the angle or distance between the two optical parts from one session to another can affect signal amplitude, contributing to inter-session variability and making it difficult to decouple positioning-induced signal changes from true variations of the machine output. Nevertheless, despite the fiber being repositioned between sessions, variability measured remained comparable to that observed with films, albeit with a slightly different calibration slope. This highlights the importance of performing a dedicated QA in the days prior to biological experiments, after the irradiation platform has been mounted. That way, reference measurements account for mounting error and eliminate this uncertainty. Besides, one of the main advantages of using this proposed setup is the accessibility and the low cost of the system compared to commercially available scintillator setups. Additionally, in our current setup the fiber was cross-calibrated against film measurements, providing relative dose output only rather than absolute dose measurements. Furthermore, recent studies have reported variations in the response of Gafchromic films to UHDR irradiations when the dose-rate is above 200-1000 Gy/s (Villoing *et al* 2022, Guan *et al* 2023, Yasuda *et al* 2026, Horst *et al* 2026). However, given the dose-rates (<100 Gy/s) used in this study, any influence on the film output is expected to be small. Also, while red-channel analysis was used due to its high sensitivity and widespread use in the radiochromic film dosimetry (Howard *et al* 2020), multichannel approaches may further reduce film- and scanner-related artifacts (Micke *et al* 2011, Darafsheh and Ghaznavi 2025). Finally, the robustness of the QA workflow was only validated using the bottom tube due to technical limitations. For a more robust validation, the QA workflow should also be validated using both tubes.

## Conclusion

In this work, we developed and validated a practical and robust dosimetry framework combining Gafchromic films and a plastic scintillating fiber for preclinical FLASH experiments using the FLASH-SARRP. As previously reported, our results show that the FLASH-SARRP system delivers dose-rates and irradiation field homogeneity compatible with preclinical FLASH studies.

Importantly, our study revealed large inter-session variability and tube asynchronicity. While the latter is intrinsic to the irradiator and can only be recorded to provide a more comprehensive understanding of the biological outcomes, the former represents a major source of experimental uncertainty that can compromise reproducibility if not properly addressed.

We demonstrated that this variability was effectively mitigated using the proposed QA workflow for FLASH preclinical experiments, integrating real-time fiber monitoring with film-based dosimetry. This approach improves the reliability and reproducibility of FLASH irradiations by correcting day-to-day output fluctuations, providing a practical framework for standardized preclinical studies.

## Fundings

Swiss National Science Foundation REquip grant IZSEZ0_232564 as well as equipment support from the University of Geneva, Geneva University Hospital and Société Académique de Genève (SACAD) to PT and MCV for the acquisition of the FLASH-SARRP.

National Institutes of Health grant R01CA254892-1 to MCV supporting MK. DFG grant to JL.

The authors are thankful for Luxium solutions for providing the scintillating fiber used in this study.

## References

Baikalov A, Tho D, Liu K, Bartzsch S, Beddar S and Schüler E 2025 Characterization of a Time-Resolved, Real-Time Scintillation Dosimetry System for Ultra-High Dose-Rate Radiation Therapy Applications Int. J. Radiat. Oncol. Biol. Phys. 121 1372–83

Böhlen T, Psoroulas S, Aylward J D, Beddar S, Douralis A, Delpon G, Garibaldi C, Gasparini A, Schüler E, Stephan F, Moeckli R and Subiel A 2024 Recording and reporting of ultra-high dose rate “FLASH” delivery for preclinical and clinical settings Radiotherapy and Oncology 200 Online: https://pubmed.ncbi.nlm.nih.gov/39245070/

Brown K H, Ghita-Pettigrew M, McIvor M P, McDowell M P, McLaughlin O, Prise K M, Sforza D, Wong J W, Rezaee M, McMahon S J and Butterworth K T 2025 Dose, dose rate and split dose impacts murine skin responses following photon FLASH irradiation Radiotherapy and Oncology 212 111125 Online: https://www.sciencedirect.com/science/article/pii/S0167814025046298?ref=pdf_download&fr=RR-2&rr=9838644119336aa0

Buranurak S, Andersen C E, Beierholm A R and Lindvold L R 2013 Temperature variations as a source of uncertainty in medical fiber-coupled organic plastic scintillator dosimetry Radiat. Meas. 56 307–11 Online: 10.1063/1.3576161

Darafsheh A and Ghaznavi H 2025 A review on radiochromic film dosimetry in radiation therapy J. Appl. Clin. Med. Phys. 26

Desrosiers M, DeWerd L, Deye J, Lindsay P, Murphy M K, Mitch M, Macchiarini F, Stojadinovic S and Stone H 2013 The Importance of Dosimetry Standardization in Radiobiology J. Res. Natl. Inst. Stand. Technol. 118 403 Online: https://pubmed.ncbi.nlm.nih.gov/26401441/

Favaudon V, Caplier L, Monceau V, Pouzoulet F, Sayarath M, Fouillade C, Poupon M F, Brito I, Hupé P, Bourhis J, Hall J, Fontaine J J and Vozenin M C 2014 Ultrahigh dose-rate FLASH irradiation increases the differential response between normal and tumor tissue in mice Sci. Transl. Med. 6 Online: 10.1126/scitranslmed.3008973

Feddersen T V, Rowshanfarzad P, Abel T N and Ebert M A 2019 Commissioning and performance characteristics of a pre-clinical image-guided radiotherapy system 42 541–51 Online: 10.1007/s13246-019-00755-4

Gao F, Yang Y, Zhu H, Wang J, Xiao D, Zhou Z, Dai T, Zhang Y, Feng G, Li J, Lin B, Xie G, Ke Q, Zhou K, Li P, Shen X, Wang H, Yan L, Lao C, Shan L, Li M, Lu Y, Chen M, Feng S, Zhao J, Wu D and Du X 2022 First demonstration of the FLASH effect with ultrahigh dose rate high-energy X-rays Radiotherapy and Oncology 166 44–50

Ghita-Pettigrew M, Brown K H, Kerr B N, Walls G M, Verginadis I I, Adrian G, Petersson K, McMahon S J and Butterworth K T 2026 Photon FLASH spares radiation-induced changes in cardiac function, remodelling and arrythmia in a preclinical model Radiotherapy and Oncology 216 111369

Guan F, Wang X, Yang M, Draeger E, Han D, Iga K, Guo F, Perles L, Li Y, Sahoo N, Mohan R and Chen Z 2023 Dosimetric response of Gafchromic^TM^ EBT-XD film to therapeutic protons Precis. Radiat. Oncol. 7 15–26 Online: 10.1002/pro6.1187

Hao X, Du H, Lin B, Wang D, Wu W, Tang M, Zhang H, Zhu Y, Zhang Y, Yang Y and Du X 2026a Preliminary mechanistic study of mitochondrial function in intestinal protection mediated by high-energy X-ray FLASH radiotherapy Radiation Oncology 2026 21:1 21 47- Online: https://link.springer.com/article/10.1186/s13014-026-02809-w

Hao X, Du H, Lin B, Wang D, Wu W, Tang M, Zhang H, Zhu Y, Zhang Y, Yang Y and Du X 2026b Preliminary mechanistic study of mitochondrial function in intestinal protection mediated by high-energy X-ray FLASH radiotherapy Radiat. Oncol. 21 Online: https://pubmed.ncbi.nlm.nih.gov/41764525/

Hart A, Cecchi D, Giguère C, Larose F, Therriault-Proulx F, Esplen N, Beaulieu L and Bazalova-Carter M 2022 Lead-doped scintillator dosimeters for detection of ultrahigh dose-rate x-rays Phys. Med. Biol. 67 Online: https://pubmed.ncbi.nlm.nih.gov/35453128/

Hill M A, Silvestre Patallo I, Aitkenhead A H, Bazalova-Carter M, Carter R, Nill S, Nisbet A, Ghita-Pettigrew M, Poirier Y, Prise K M, Subiel A and Schettino G 2025 The importance of standardization and challenges of dosimetry in conventional preclinical radiation biology research Br. J. Radiol. 98 993–1004 Online: https://pubmed.ncbi.nlm.nih.gov/40127177/

Horst F, Schüller A, Kapsch R-P, Hackel T, Pehlivan J, Schott C, Beyreuther E and Pawelke J 2026 Physics in Medicine & Biology OPEN ACCESS RECEIVED Study of the over-response of radiochromic films in ultra-high dose rate and dose per pulse electron beams Phys. Med. Biol 71 45020 Online: 10.1088/1361-6560/ae47b6

Howard M E, Herman M G and Grams M P 2020 Methodology for radiochromic film analysis using FilmQA Pro and ImageJ PLoS One 15 e0233562 Online: https://pmc.ncbi.nlm.nih.gov/articles/PMC7241712/

Jaccard M, Durán M T, Petersson K, Germond J F, Liger P, Vozenin M C, Bourhis J, Bochud F and Bailat C 2018 High dose-per-pulse electron beam dosimetry: Commissioning of the Oriatron eRT6 prototype linear accelerator for preclinical use: Commissioning *Med*. Phys. 45 863–74

Jorge P G, Jaccard M, Petersson K, Gondré M, Durán M T, Desorgher L, Germond J F, Liger P, Vozenin M C, Bourhis J, Bochud F, Moeckli R and Bailat C 2019 Dosimetric and preparation procedures for irradiating biological models with pulsed electron beam at ultra-high dose-rate Radiotherapy and Oncology 139 34–9

Kirsch D G, Diehn M, Kesarwala A H, Maity A, Morgan M A, Schwarz J K, Bristow R, Demaria S, Eke I, Griffin R J, Haas-Kogan D, Higgins G S, Kimmelman A C, Kimple R J, Lombaert I M, Ma L, Marples B, Pajonk F, Park C C, Schaue D and Bernhard E J 2018 The Future of Radiobiology J. Natl. Cancer Inst. 110 Online: https://pubmed.ncbi.nlm.nih.gov/29126306/

Knol M, Kunz L, Ollivier J, Limoli C, Tsoutsou P and Vozenin M-C 2026 In Regard to Grilj et al International Journal of Radiation Oncology*Biology*Physics 125 665–7 Online: https://linkinghub.elsevier.com/retrieve/pii/S0360301626003974

Lin B, Du H, Yang Y, Hao X, Gao F, Liang Y, Tang W, Xu H, Tang M, Liao Y, Wang D, Lin B, Zhu Y, Wang T, Gu R, Mao X, He Y, Zhang Y, Li J, Zhou Z, Wang J, Wu D and Du X 2025 FLASH Radiation Therapy Using High-Energy x Rays: Validation of the Flash Effect Triggered by a Compact Device Int. J. Radiat. Oncol. Biol. Phys.

Lin B, Du H, Yang Y, Hao X, Gao F, Liang Y, Tang W, Xu H, Tang M, Liao Y, Wang D, Lin B, Zhu Y, Wang T, Gu R, Mao X, He Y, Zhang Y, Li J, Zhou Z, Wang J, Wu D and Du X 2026 FLASH Radiation Therapy Using High-Energy X-Rays: Validation of the Flash Effect Triggered by a Compact Device Int. J. Radiat. Oncol. Biol. Phys. 124 1084–95 Online: https://pubmed.ncbi.nlm.nih.gov/41093135/

Liu K, Waldrop T, Aguilar E, Mims N, Neill D, Delahoussaye A, Li Z, Swanson D, Lin S H, Koong A C, Taniguchi C M, Loo B W, Mitra D and Schüler E 2025 Redefining FLASH Radiation Therapy: The Impact of Mean Dose Rate and Dose Per Pulse in the Gastrointestinal Tract International Journal of Radiation Oncology*Biology*Physics 121 1063–76 Online: https://linkinghub.elsevier.com/retrieve/pii/S0360301624034667

Maxim P G, Tantawi S G and Loo B W 2019 PHASER: A platform for clinical translation of FLASH cancer radiotherapy Radiotherapy and Oncology 139 28–33 Online: https://pubmed.ncbi.nlm.nih.gov/31178058/

Micke A, Lewis D F and Yu X 2011 Multichannel film dosimetry with nonuniformity correction Med. Phys. 38 2523–34 Online: https://pubmed.ncbi.nlm.nih.gov/21776787/

Miles D, Sforza D, Wong J and Rezaee M 2024 Dosimetric characterization of a rotating anode x-ray tube for FLASH radiotherapy research Med. Phys. 51 1474–83 Online: https://pubmed.ncbi.nlm.nih.gov/37458068/

Miles D, Sforza D, Wong J W, Gabrielson K, Aziz K, Mahesh M, Coulter J B, Siddiqui I, Tran P T, Viswanathan A N and Rezaee M 2023 FLASH Effects Induced by Orthovoltage X-Rays Int. J. Radiat. Oncol. Biol. Phys. 117 1018 Online: https://pmc.ncbi.nlm.nih.gov/articles/PMC11189000/

Montay-Gruel P, Bouchet A, Jaccard M, Patin D, Serduc R, Aim W, Petersson K, Petit B, Bailat C, Bourhis J, Bräuer-Krisch E and Vozenin M C 2018 X-rays can trigger the FLASH effect: Ultra-high dose-rate synchrotron light source prevents normal brain injury after whole brain irradiation in mice Radiotherapy and Oncology 129 582–8 Online: https://pubmed.ncbi.nlm.nih.gov/30177374/

Moreau M, Kelly K, Mao S, China D, Wasihun G, Pandya A, Tajik-Mansoury M A, Sforza D, Miles D, Narang A K, Rezaee M, Ngwa W and Ding K 2025 Leveraging the Immune Response from LIFE Biomaterial and Photon-Flash in Pre-Clinical Pancreatic Cancer Treatment Pharmaceutics 17

Rezaee M, Iordachita I and Wong J W 2021 Ultrahigh dose-rate (FLASH) x-ray irradiator for pre-clinical laboratory research Phys. Med. Biol 66 95006 Online: 10.1088/1361-6560/abf2fa

Tajik Mansoury M A, Sforza D, Wong J, Iordachita I and Rezaee M 2025 Dosimetric commissioning of small animal FLASH radiation research platform Phys. Med. Biol. 70

Villoing D, Koumeir C, Bongrand A, Guertin A, Haddad F, Métivier V, Poirier F, Potiron V, Servagent N, Supiot S, Delpon G and Chiavassa S 2022 Technical note: Proton beam dosimetry at ultra-high dose rates (FLASH): Evaluation of GAFchromic^TM^ (EBT3, EBT-XD) and OrthoChromic (OC-1) film performances Med. Phys. 49 2732–45

Vozenin M C, Bourhis J and Durante M 2022 Towards clinical translation of FLASH radiotherapy Nat. Rev. Clin. Oncol. 19 791–803

Vozenin M C, Loo B W, Tantawi S, Maxim P G, Spitz D R, Bailat C and Limoli C L 2024 FLASH: New intersection of physics, chemistry, biology, and cancer medicine Rev. Mod. Phys. 96

Vozenin M C, Montay-Gruel P, Limoli C and Germond J F 2020 All Irradiations that are Ultra-High Dose Rate may not be FLASH: The Critical Importance of Beam Parameter Characterization and In Vivo Validation of the FLASH Effect Radiat. Res. 194 571–2 Online: https://pubmed.ncbi.nlm.nih.gov/32853355/

Vozenin M C, Montay-Gruel P, Tsoutsou P and Limoli C L 2025 Mechanisms, challenges and opportunities for FLASH radiotherapy in cancer Nature Reviews Cancer 2025 26:1 26 62–75 Online: https://www.nature.com/articles/s41568-025-00878-9

Wang Y F, Lin S C, Na Y H, Black P J and Wuu C S 2017 Dosimetric Verification and Commissioning for an Image-Guided Small Animal Irradiator Int. J. Radiat. Oncol. Biol. Phys. 99 E624

Yasuda H, Toshito T, Umezawa M, Yamada M, Tanaka K, Omachi C, Yogo K, Bantan H and Nakashima T 2026 Comparative analysis of the responses of EBT-XD and EBT4 films to ultra-high dose rate proton beams Radiation Physics and Chemistry 239 113349 Online: https://www.sciencedirect.com/science/article/pii/S0969806X25008412?via%3Dihub

Zhang S, Peng Q, Zhang J, Wang Z, Cheng X, Zhang Y and Cao Z 2026 Characterization of the Intestinal Metabolic Profiles Following Photon FLASH Irradiation Dose-Response 24

Zhu H, Liu S, Qiu J, Hu A, Zhou W, Wang J, Gu W, Zhu Y, Zha H, Xiang R, Li J, Qiu R, Zhao C, Huang P and Deng X 2025 Instantaneous dose rate as a crucial factor in reducing mortality and normal tissue toxicities in murine total-body irradiation: a comparative study of dose rate combinations Molecular Medicine 31

