## Supplementary material for "Dosimetric Characterization and Workflow Optimization of the FLASH-SARRP for Reliable Preclinical Radiobiological Studies": supp data

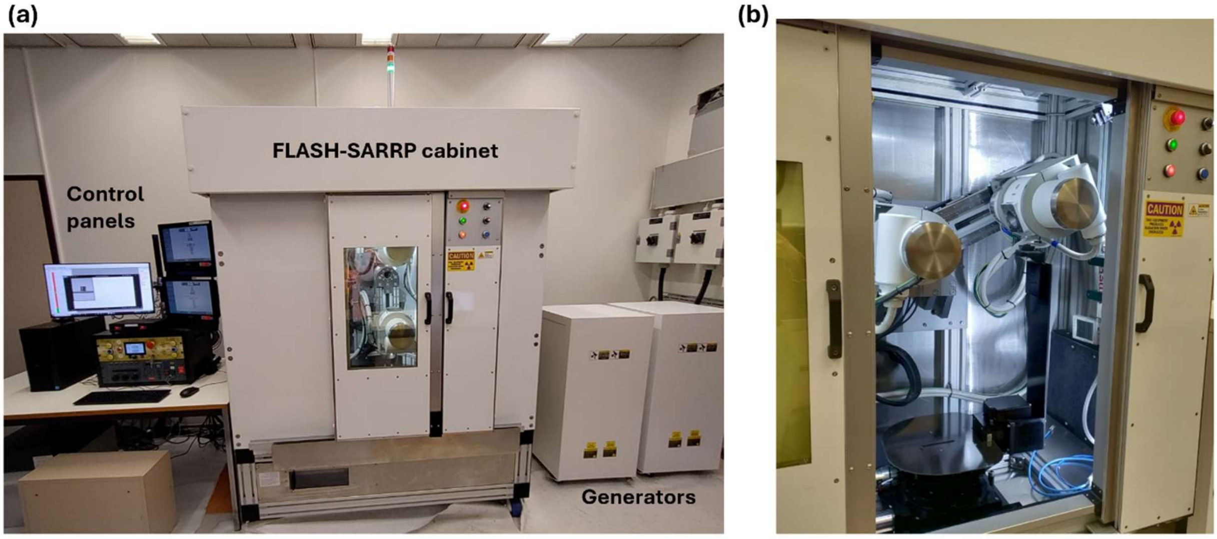


**Supplementary Figure 1. Overview of the FLASH-SARRP irradiation platform. (a)** External view of the FLASH-SARRP system, including the irradiation cabinet, control electronics, and the generators. **(b)** Interior of the irradiation cabinet showing the movable dual-tube X-ray configuration.

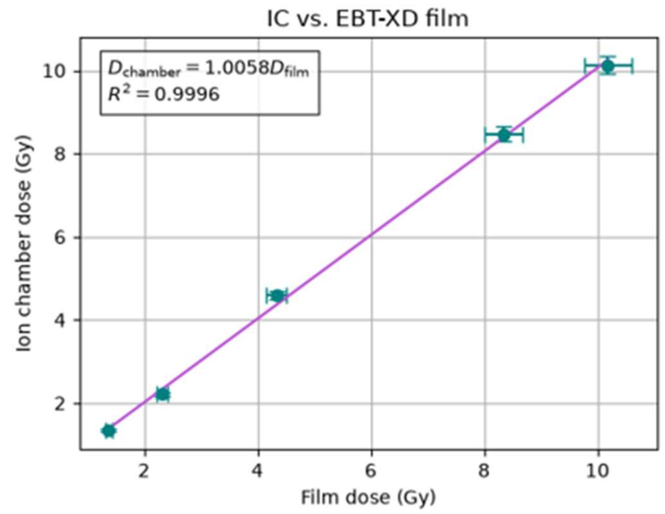


**Supplementary Figure 2. Validation of EBT-XD film dosimetry using an ionization chamber. (a)** Comparison between doses measured with EBT-XD film and a calibrated ionization chamber (IC) under identical irradiation conditions (n=3 for IC measurements, n=2 for film measurements, mean is plotted with 4% errorbars for the film measurements and 2.1% for the IC chamber). 150 kV, 630 mA, and varying exposure times between 32 and 250 ms were used. A linear regression constrained through the origin was fitted with a slope of 1.0058 and $R^2$ = 0.9996. The observed deviation from unity (0.58%) was smaller than the uncertainty related to the film analysis (4%).

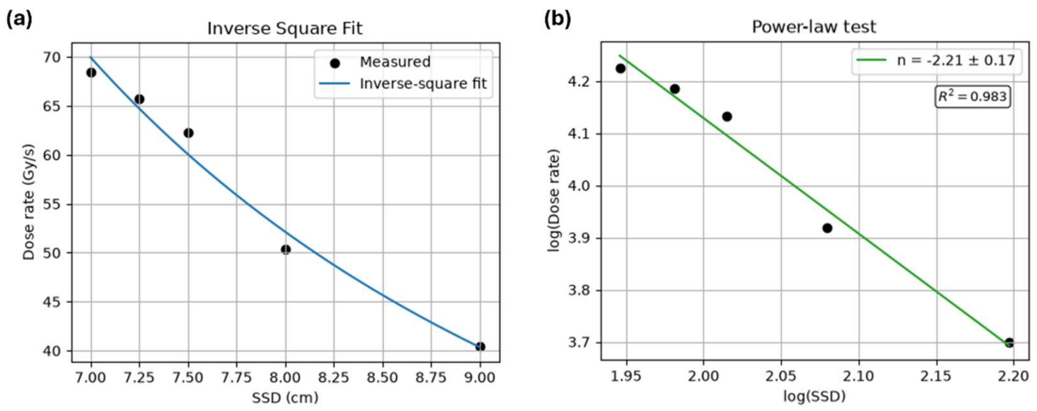


**Supplementary Figure 3. Experimental validation of the inverse square approximation. (a)** Measured dose-rate as a function of the source-to-surface distance (SSD) for irradiations performed at 150 kV, 630 mA, and 125 ms. An inverse square model $dose\ rate = \frac{k}{(SSD+d_0)^2}$ , with a potential offset $d_0$, was fitted through the measurements, resulting in $k = 2.79 \cdot 10^3$ and $d_0 = -0.69 \pm 0.67\ cm$. The individual points represent single film measurements with an overall uncertainty of the film analysis of 4%. **(b)** Power-law analysis of the measured dose-rate dependence on the SSD. The fitted exponent of -2.21 ± 0.17 is consistent within experimental uncertainty with the ideal inverse-square dependence of n = -2.

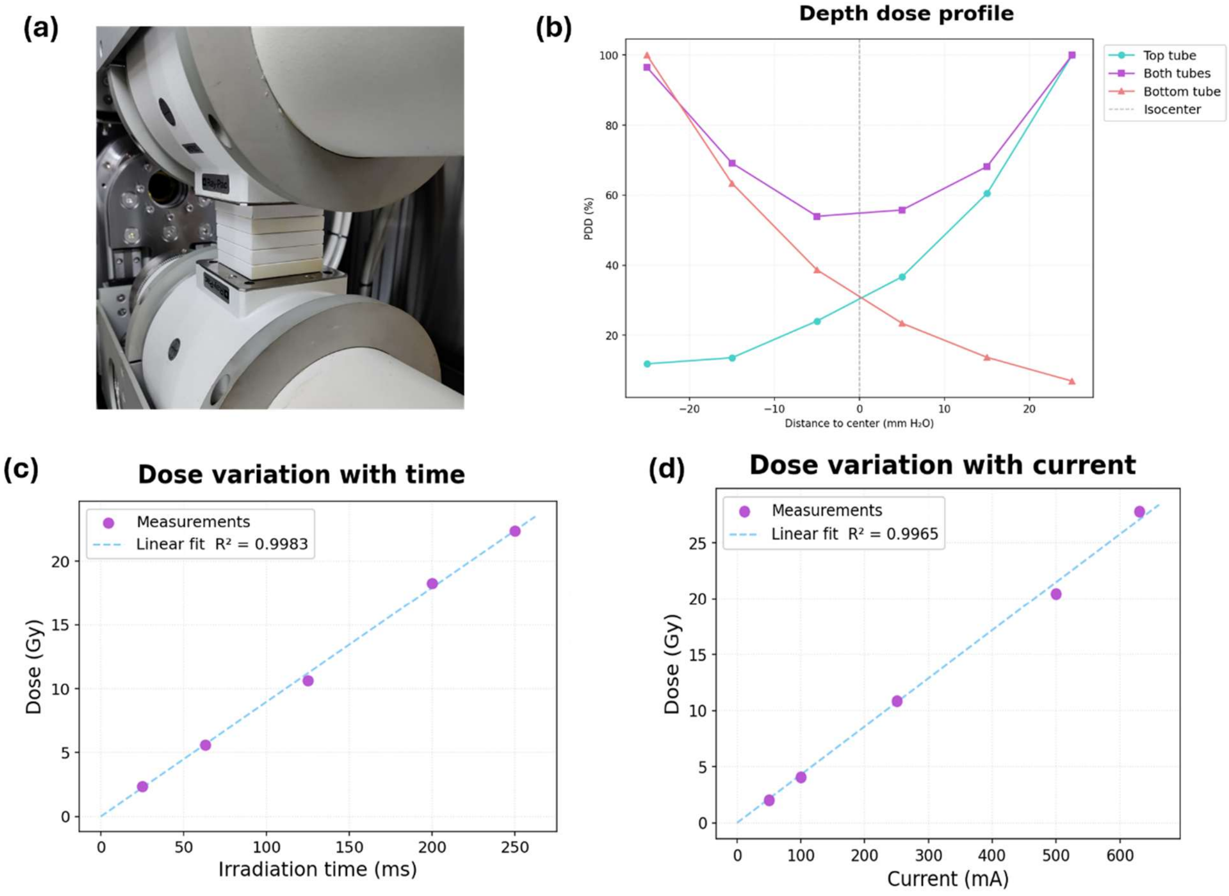


**Supplementary Figure 4. Beam Characteristics. (a)** A photo of the setup used to obtain the PDD. Gafchromic film was placed between the solid water slabs, as well as directly at the surface of both tubes. **(b)** Percentage Depth Dose (PDD) curves of irradiations performed with the top (blue), bottom (red), and both tubes (purple) in solid water slabs. **(c)** Linearity of the irradiation time, measured with Gafchromic films at SSD = 7 cm from both tubes. A linear fit is determined based on these measurements. **(d)** Linearity of the current, measured with Gafchromic film at SSD = 7 cm from both tubes.

| Session | Date | Dosimeter(s) | Tubes | Figure |
|---|---|---|---|---|
| S1 | 08/Jul/25 | Film | Both | 4c, d, e, f |
| S2 | 15/Jul/25 | Film | Both | 4c, d, e, f |
| S3 | 22/Jul/25 | Film | Both | 4c, d, e, f |
| S4 | 06/Sep/25 | Film | Both | 4e, f |
| S5 | 16/Feb/26 | Film, Fiber | Both | 3d |
| S6 | 17/Feb/26 | Film, Fiber | Both | 3d |
| S7 | 18/Feb/26 | Film, Fiber | Both | 3d |
| S8 | 24/Feb/2026 (morning) | Film, Fiber | Bottom | 6b, c |
| S9 | 24/Feb/2026 (afternoon) | Film, Fiber | Bottom | 6b, c |
| S10 | 02/Mar/26 | Fiber | Bottom | 3a |
| S11 | 09/Mar/26 | Film, Fiber | Bottom | 6b, c |
| S12 | 12/Mar/26 | Film, Fiber | Bottom | 6b, c |
| S13 | 11/May/26 | Film | Both | 4a, b |
| S14 | 12/May/26 | Film | Both | 4a, b |
| S15 | 20/May/26 | Fiber | Both | 5b; 3b |
| S16 | 26/May/26 | Film, Fiber | Both | 5b; 3d; 4a, b |
| S17 | 27/May/26 | Film, Fiber | Both | 5a, b; 4a, b |
| S18 | 28/May/26 | Film | Both | 4a, b |
| S19 | 29/May/26 | Fiber | Both | 5a, b |
| S20 | 01/Jun/26 | Film, Fiber | Both | 3d; 4a, b |
| S21 | 02/Jun/26 | Film | Both | 4a, b |
| S22 | 03/Jun/26 | Film | Both | 4a, b |

**Supplementary Table 1. Overview of all experimental sessions.**
